# NAD^+^ repletion with niacin counteracts cancer cachexia

**DOI:** 10.1101/2022.07.06.499010

**Authors:** Marc Beltrà, Noora Pöllänen, Claudia Fornelli, Kialiina Tonttila, Myriam Y. Hsu, Sandra Zampieri, Lucia Moletta, Paolo E. Porporato, Riikka Kivelä, Marco Sandri, Juha J. Hulmi, Roberta Sartori, Eija Pirinen, Fabio Penna

## Abstract

Cachexia is a debilitating wasting syndrome and highly prevalent comorbidity in cancer patients. It manifests especially with energy and mitochondrial metabolism aberrations that promote tissue wasting. We recently identified nicotinamide adenine dinucleotide (NAD^+^) loss to associate with muscle mitochondrial dysfunction in cancer hosts. In this study we confirmed that depletion of NAD^+^ and downregulation of *Nrk2*, an NAD^+^ biosynthetic enzyme, are common features of different mouse models and cachectic cancer patients. Testing NAD^+^ repletion therapy in cachectic mice revealed that NAD^+^ precursor, vitamin B3 niacin, efficiently corrected tissue NAD^+^ levels, improved mitochondrial metabolism and ameliorated cancer- and chemotherapy-induced cachexia. To examine NAD^+^ metabolism in a clinical setting, we showed that the low expression of *NRK2* in cancer patients correlated with metabolic abnormalities underscoring the significance of NAD^+^ in the pathophysiology of human cancer cachexia. Overall, our results propose a novel therapy target, NAD^+^ metabolism, for cachectic cancer patients.

## INTRODUCTION

Cancer cachexia (CC) is a complex multifactorial syndrome resulting from both tumor-induced host adaptation and anti-cancer treatment side effects, being present in and worsening the outcome of more than half of all cancer patients. CC is clinically characterized by an involuntary loss of body weight mainly due to muscle wasting, with or without depletion of adipose tissue that impair patients’ quality of life and survival ^1^. Several energy metabolic abnormalities including mitochondrial dysfunction have been characterized in CC suggesting that targeting energy metabolism could be useful when designing novel anti-cachexia treatments^2,3^. Recently, our study on C26 adenocarcinoma bearing mice showed that the decline of mitochondrial oxidative phosphorylation (OXPHOS) occurred in parallel with depleted NAD^+^ levels in the skeletal muscle^4^. Given that NAD^+^ is an essential cofactor for mitochondrial energy production, alterations of its levels can affect mitochondrial homeostasis and subsequently the function of the tissue. In support of this notion, mRNA transcript levels of NAD^+^ biosynthesis genes positively correlate with the expression of genes regulating muscle mitochondrial biogenesis, muscle mass growth and muscle regeneration in mice^5,6^. Recent rodent and human studies have reported NAD^+^ depletion as a pathological hallmark for various muscle diseases including sarcopenia and mitochondrial myopathy^6–8^. NAD^+^ depletion is typically caused by impaired NAD^+^ biosynthesis, increased activities of NAD^+^ degrading enzymes, a combination of both or changes in metabolic reactions relying on NAD^+^/NADH redox couple. In our above-mentioned work in C26-bearing mice, skeletal muscle NAD^+^ loss was associated with a strong transcriptional downregulation of the NAD^+^ biosynthetic enzyme *nicotinamide riboside kinase 2* (*Nrk2*). This salvage pathway enzyme metabolizes vitamin B3, nicotinamide riboside (NR), towards NAD^+^ and is regulated by stress and alterations in the intracellular NAD^+^ and energy supply^9^. NR has been previously published to alleviate cachexia in mice bearing the C26 tumor^10^, although the demonstration of NAD^+^ replenishment is lacking. The clinical use of NR is challenged as the clinical trials published so far have failed to demonstrate positive outcomes on tissue energy metabolism^11,12^. In contrast, the other vitamin B3 form, niacin (NA), has a proven safety record in humans and it has been published to improve muscle mitochondrial and energy metabolism in patients with adult-onset mitochondrial myopathy^13^. Unsolved questions remain: 1) how common muscle NAD^+^ depletion and *Nrk2* downregulation are in CC induced by distinct tumors, 2) is NAD^+^ metabolism disturbed in other tissues beyond the skeletal muscle, and 3) can NAD^+^ repletion with NA mitigate the symptoms of cancer cachexia.

The current study aimed to better characterize NAD^+^ and energy metabolism impairments in CC and examine the potential therapeutic role of NA supplementation. Skeletal muscle NAD^+^ deficiency was detected in mice with severe CC, whereas muscle *Nrk2* loss was observed in several preclinical CC models and validated in cancer patient muscle biopsies. In addition, the depletion of all NAD metabolites was observed in the liver of mice suffering from acute or chronic CC. NA corrected NAD^+^ deficiency and increased mitochondrial biogenesis in both skeletal muscle and liver of cachectic mice, partially restoring muscle mass loss and energy metabolism changes. Thus, our findings propose that *Nrk2* repression and NAD^+^ metabolism aberrations are prevalent features of CC. Correcting NAD^+^ metabolism has a protective role in maintaining adequate energy homeostasis and preventing cachexia in tumor-bearing animals.

## RESULTS

### Impaired skeletal muscle NAD^+^ metabolism is a common feature of experimental cancer cachexia

In this study, an exacerbation of muscle NAD^+^ depletion was observed in C26-bearing animals treated with Folfox chemotherapy (C26-F, Fig. 1a) in comparison to our previous publication in chemotherapy-naive C26-mice (NAD^+^ decrease: 30,8% versus 12,5%)^4^. Consistently, a reduction in NAD^+^ was observed also in the skeletal muscle of KPC tumor-bearing mice (Fig. 1a), a CC model representative of pancreatic ductal adenocarcinoma^14^. In order to better model CC, we assessed NAD^+^ content in the skeletal muscle of Villin-Cre/Msh2^loxP/loxP^ (VCM) mice that slowly and spontaneously develop neoplasms due to the conditional knock-out of the mismatch repair gene Msh2 in the enterocytes of the intestinal mucosa^15^ (a characterization of this new cachexia model is provided in Supplementary Fig. 1). Although presenting with significant anemia, body weight loss and muscle wasting due to tumor progression is moderate in the VCM model compared to aged-matched *Msh2*^loxP/loxP^ mice (controls; Supplementary Fig. 1a-c), supporting the idea of a milder and more chronic model of CC. As opposed to the previous severe models of CC, skeletal muscle NAD^+^ levels were not significantly reduced in VCM mice (Fig. 1a).

**Figure 1.**
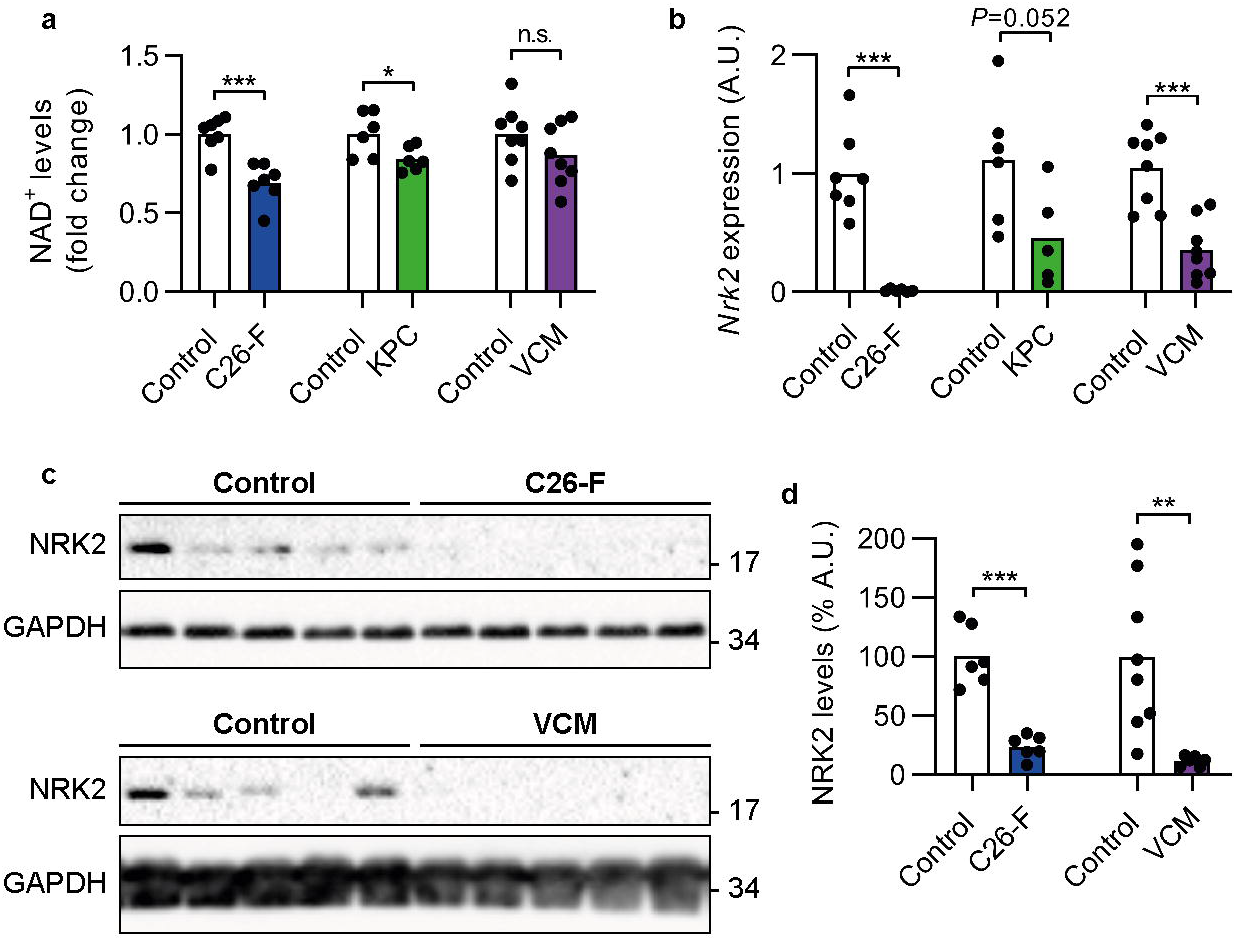
NAD^+^ depletion and Nrk2 downregulation as hallmarks of cancer cachexia in the skeletal muscle. **a** NAD^+^ levels in the skeletal muscle of C26-F (*n*=7), KPC (*n*=6) and VCM CC models (*n*=8). **b** RT-qPCR quantification of *Nrk2* gene expression in C26-F (*n*=6-7), KPC (*n*=5-6) and VCM (*n*=8) mice. Data are normalized to housekeeping gene expression and displayed as relative expression. **c, d** Representative western blotting bands and densitometry analysis of NRK2 protein levels on C26-F (*n*=6) and VCM (*n*=6-8) mice. GAPDH expression was used as loading control. Data are shown as means with individual values. Statistical analysis was performed using Student’s t-test, \**P*<0.05, \*\**P*<0.01, \*\*\**P*<0.001. and n.s.: non-significant. A.U.; arbitrary units.

In chemotherapy-naive C26 tumor-bearing mice, decreased NAD^+^ levels associated with the repression of the NAD^+^ biosynthetic gene *Nrk2*^4^. Consistently, cachectic mice showed downregulation of muscle *Nrk2* gene expression in both C26-F and VCM models compared to controls, while a trend towards *Nrk2* reduction was detected in KPC mice (Fig. 1b). Remarkably, NRK2 protein levels were nearly undetectable in muscle homogenates of C26-F and VCM mice when compared to their respective control group (Fig. 1c-d). Altogether, these data demonstrate that skeletal muscle NAD^+^ content is depleted in severe CC and that *Nrk2* downregulation is a common feature of experimental CC independently from the severity of the syndrome.

### Muscle *NRK2* loss occurs in human cancer cachexia and metabolome profiling reveals a unique signature of low *NRK2*-expressing skeletal muscle

To validate the preclinical data in humans, *NRK2* expression was assessed in skeletal muscle biopsies from colorectal and pancreatic cancer patients and compared to healthy subjects. The samples originate, with some additions, from a previous study^16^ (patient characteristics are summarized in Supplementary Table 1). *NRK2* expression decreased in patients classified as pre-cachectic and showed a further pronounced downregulation in cachectic patients (Fig. 2a) confirming that *NRK2* loss is a novel common alteration in CC. To examine the relationship between *NRK2* loss and muscle metabolism in CC, we selected 10 patients with the highest (comparable to healthy controls) and 10 with the lowest (almost 10-fold decrease) *NRK2* expression that did not differ in terms of body weight loss (Fig. 2b-c). Moreover, *NRK2* expression levels were independent from muscle mass and wasting, as no association was found with macroscopic clinical features of the current cachexia diagnostic criteria, mainly based on body weight loss and sarcopenia (Supplementary Table 2-3). A metabolomic characterization revealed a peculiar signature in the low *NRK2* expressing muscles clearly distinguished from high *NRK2* and healthy counterparts (Fig. 2d). Low *NRK2* samples showed an accumulation of glycolysis intermediates, nucleotides and amino acids, indicative of impaired energy metabolism and hypercatabolic state (Fig. 2e-h). This trait can be partially observed also in the sera of the same individuals (Fig. 2i,j), suggesting that muscle energy failure and protein hypercatabolism could be diagnosed with minimally invasive procedures.

**Figure 2.**
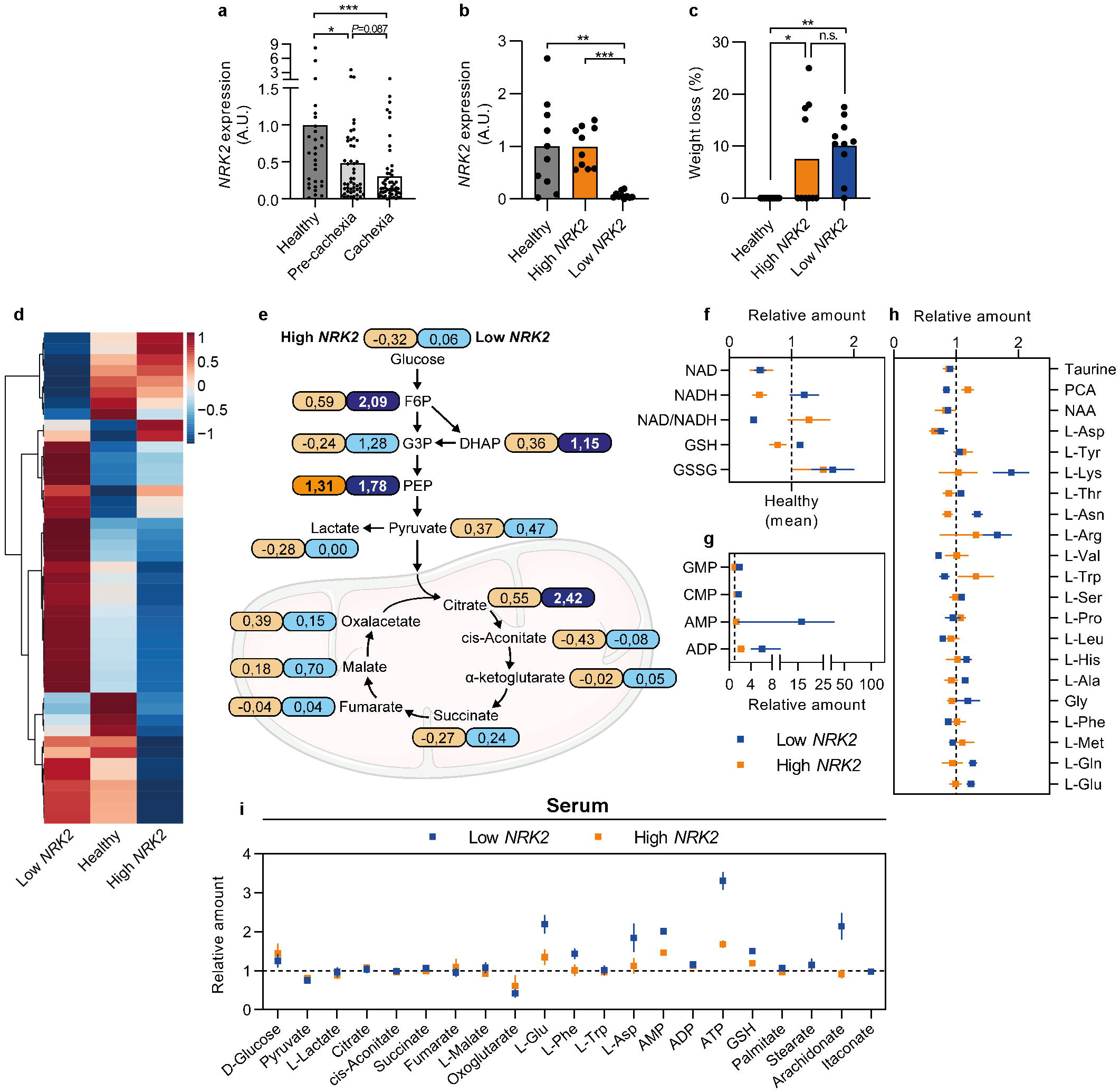
Metabolome analysis in human muscle biopsies and serum according to *NRK2* expression. **a** Relative expression of *NRK2* transcript in *rectus abdominis* muscle biopsies from healthy volunteers (*n*=28) and cancer patients stratified in pre-cachectic (*n*=49) and cachectic (*n*=53) assessed by RT-qPCR. **b** Relative gene expression of *NRK2* in healthy volunteers (*n*=10; grey), high *NRK2*-expressing (*n*=10; orange) and low *NRK2*-expressing (*n*=10; blue) groups. **c** Weight loss percentage in the previous 6 months before biopsy collection from patients with high or low *NRK2* expression that were selected for metabolome analysis. Statistical analysis was performed using Kruskal-Wallis test with adjustment for multiple testing (Benjamini, Krieger and Yekuteli): \**P*<0.05, \*\**P*<0.01, \*\*\**P*<0.001 and n.s.: non-significant. **d** Heatmap highlighting the abundancy of all the detected metabolites in muscle biopsies. Color represents the mean Z score of each group. **e** Schematic illustration presenting the relative abundance of metabolites categorized in the “glycolysis” and the “Kreb’s Cycle” pathways. Numbers in boxes represent fold change of either high *NRK2* (orange) or low *NRK2* (blue) groups compared to healthy volunteers for each metabolite. **f-h** Relative abundance of metabolites detected in muscle samples categorized as (**f**) “redox”, (**g**) “nucleotides”, and (**h**) “amino acids”. **i** Relative abundance of detectable metabolites in the sera. Data display: **a**-**c** are means with individual values, **e** are fold change *vs* Healthy group, **f-i** are relative abundance ± SEM *vs* Healthy group (dotted line). PCA; pyroglutamate, NAA; N-acetyl aspartate, A.U.; arbitrary units.

### Niacin rescues skeletal muscle NAD^+^ levels, ameliorates muscle wasting and improves protein metabolism in experimental cancer cachexia

NRKs catalyze the utilization of NAD^+^ precursor NR via the salvage pathway. In the skeletal muscle, *Nrk2* is the most expressed isoform in both BALB/c and C57BL/6 strains as compared to *Nrk1* (Supplementary Fig. 2a). Considering the *Nrk2* repression in CC and the lack of a simple, translatable tool to correct *Nrk2* expression, we decided to use the NAD^+^ booster NA, a precursor that is utilized for NAD^+^ biosynthesis through the Preiss-Handler pathway thus bypassing NRK2^17^. C26-F animals were treated with a daily dose (150 mg/kg) of NA starting from day 4 after C26 implantation until day 28 (Fig. 3a). In addition to NAD^+^ depletion (Fig. 1a), C26-F mice presented with a significant decrease of NADH and NADPH levels while NADP levels were similar in comparison to controls (Supplementary Fig. 2b). Besides *Nrk2 l*oss, C26-F mice showed an overall repression of genes involved in NAD^+^ biosynthesis via the salvage and Preiss-Handler pathways (Supplementary Fig. 2c) and enhanced enzyme activity of poly(ADP-ribose)polymerases (PARPs), one of the main consumers of cellular NAD^+^ pool operating for example in DNA repair (Supplementary Fig. 2d). Interestingly, NA increased skeletal muscle NAD^+^ and NADP^+^ concentrations almost to the control levels and slightly impacted on NADH and NADPH levels (Fig. 3b, Supplementary Fig. 2b). Moreover, NA supplementation improved cachexia symptoms by counteracting the loss of body weight and muscle mass and partially rescuing grasping strength (Fig. 3c-e, Supplementary Fig. 2e). When tested *in vitro*, NA increased C26 cell proliferation, but this effect was neutralized when administered together with oxaliplatin or 5-fluorouracil (Supplementary Fig. 2f). Additionally, NA increased the number of death cells and partially potentiated oxaliplatin toxicity (Supplementary Fig. 2g). *In vivo*, tumor mass was not significantly affected by NA in C26-F mice (Supplementary Fig. 2h). Consistent with our previous report^18^, C26-F mice presented with decreased skeletal muscle protein synthesis, increased ratio of the active LC3B isoform (LC3B-II; Fig. 3f-h) and AMPK phosphorylation (Fig. 3f-i), suggestive of increased autophagy and energy shortage, respectively. Interestingly, both protein synthesis and LC3B-II accumulation were in part rescued by NA (Fig. 3f-h), while AMPK activation was partially prevented (Fig. 3f,i).

**Figure 3.**
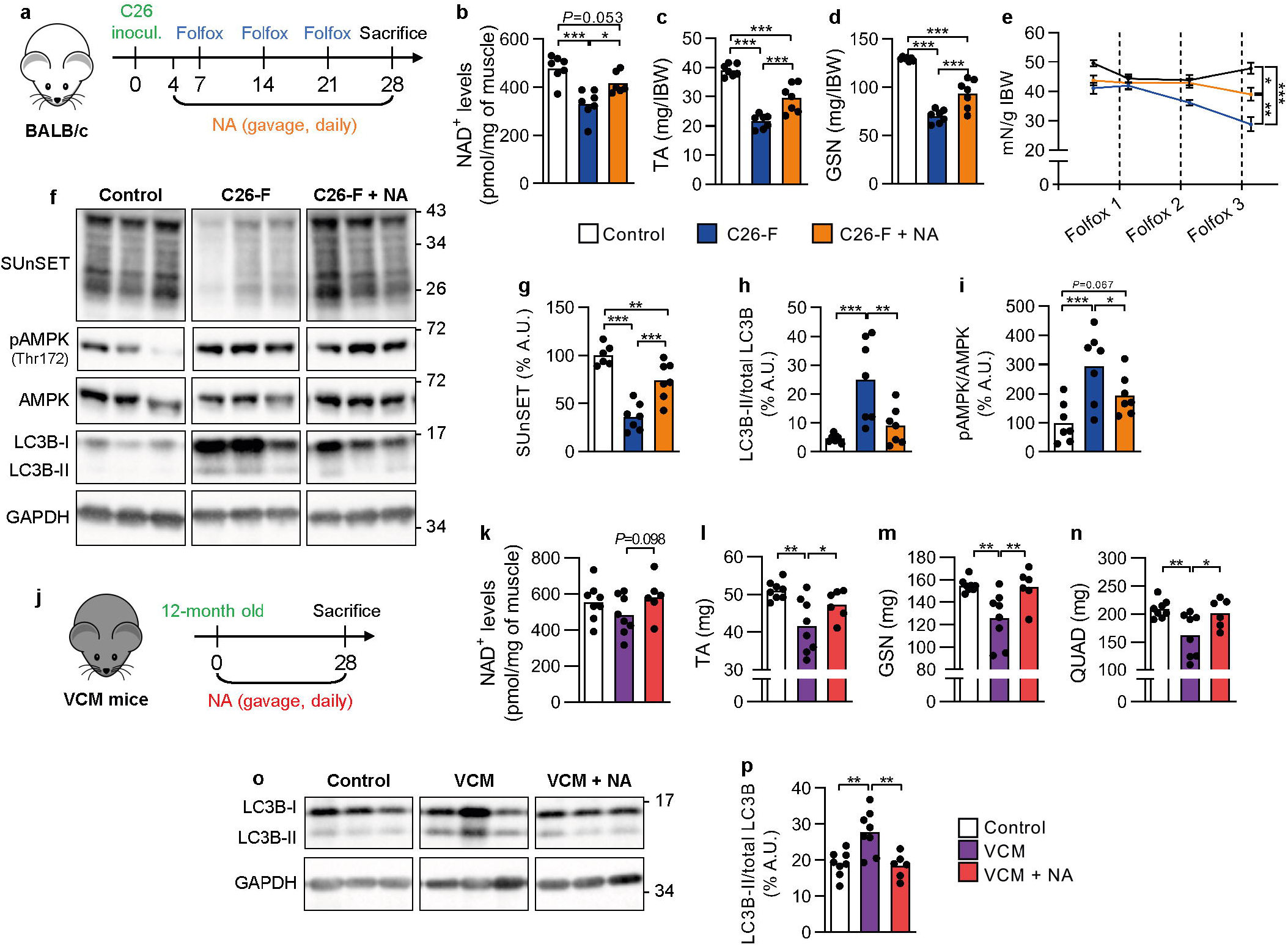
Effects of NA treatment on muscle mass, muscle function and protein metabolism in CC mouse models. **a** Study design of the C26-F model and NA treatment with three experimental groups: control (*n*=6-7), C26-F (*n*=7) and C26-F+NA (*n*=6-7). **b** NAD^+^ levels of control, C26-F and C26-F+NA groups represented as pmol/mg of muscle weight **c, d** *Tibialis anterior* (TA) and *gastrocnemius* (GSN) muscle wet weight normalized by initial body weight (IBW). **e** Grasping strength at the start of NA treatment and the day after every Folfox administration. **f-i** Representative western blotting bands (**f**) and densitometry analysis of puromycin incorporation (SUnSET analysis) (**g**), LC3B-II normalized to total LC3B (**h**) and p-AMPK normalized to total AMPK (**i**) protein levels. GAPDH protein expression was used as loading control. **j** Experimental design of the VCM model and NA treatment with three experimental groups: control (*n*=8), VCM (*n*=8) and VCM + NA (*n*=6). **k** NAD^+^ levels of control, VCM and VCM+NA groups represented as pmol/mg of muscle weight. **l-n** TA, GSN and *quadriceps femoris* (QUAD) muscle wet weight. **o, p** Representative western blotting bands (**o**) and densitometry analysis of LC3B-II normalized to total LC3B (**p**) protein levels. GAPDH protein expression was used as loading control. Data are shown in panels **b-d, g-i, k-n** and **p** as means with individual values and in panel **e** as means ± SEM (*n*=7). Statistical analysis was performed using ANOVA + Fisher’s LSD test. \**P*<0.05, \*\**P*<0.01 and \*\*\**P*<0.001. NA; niacin, A.U.; arbitrary units.

To further explore the impact of NA in a more chronic model of CC presenting with *Nrk2* loss, VCM mice were treated with NA for 28 days (Fig. 3j). In line with the preserved NAD^+^ content in VCM mice (Fig. 1a), NAD metabolites, the expression of NAD^+^ biosynthetic genes and PARP activity did not differ between control and VCM groups (Supplementary Fig. 2i-k). NA supplementation minimally impacted on skeletal muscle NAD^+^ and other NAD metabolites (Fig. 3k, Supplementary Fig. 2i). Nonetheless, NA partially protected VCM mice from muscle mass loss (Fig. 3l-n), preventing muscle autophagosome accumulation (Fig. 3o,p) and increased expression of E3 ubiquitin ligases, autophagy and mitophagy genes (Supplementary Fig. 2l-n). Overall, NA showed beneficial effects on CC by preventing muscle loss and induction of autophagy markers both in severe and mild CC models.

### Niacin improves skeletal muscle mitochondrial biogenesis in experimental cancer cachexia

Muscle mitochondrial dysfunction has gained importance in recent years as a crucial feature of CC^19^. In the current experiments, the skeletal muscle of C26-F mice presented with reduced protein levels of the main activator of mitochondrial biogenesis PGC-1α (Fig 4a, b), a decline in mitochondrial DNA (mtDNA) amount (Fig. 4c), and reduced expression of the mitochondrial mass marker, TOMM20, and several subunits of the mitochondrial oxidative phosphorylation complexes (Fig 4a, d-e). Additionally, several genes involved in mitochondrial biogenesis were downregulated (Supplementary Fig. 2o), whereas PINK1 protein, a marker of mitochondrial damage and depolarization^20^, accumulated in the skeletal muscle of C26-F mice (Fig 4a, f). NA increased the abundance of mtDNA and preserved PGC-1α, TOMM20 and OXPHOS complex subunit protein levels (Fig. 4a-e) without impacting on PINK1 accumulation (Fig 4a, f). No transcriptional induction of genes crucial for mitochondrial health and biogenesis was observed upon NA indicating (Supplementary Fig. 2o) that NA’s effect on mitochondrial metabolism is mediated via post-transcriptional mechanism.

**Figure 4.**
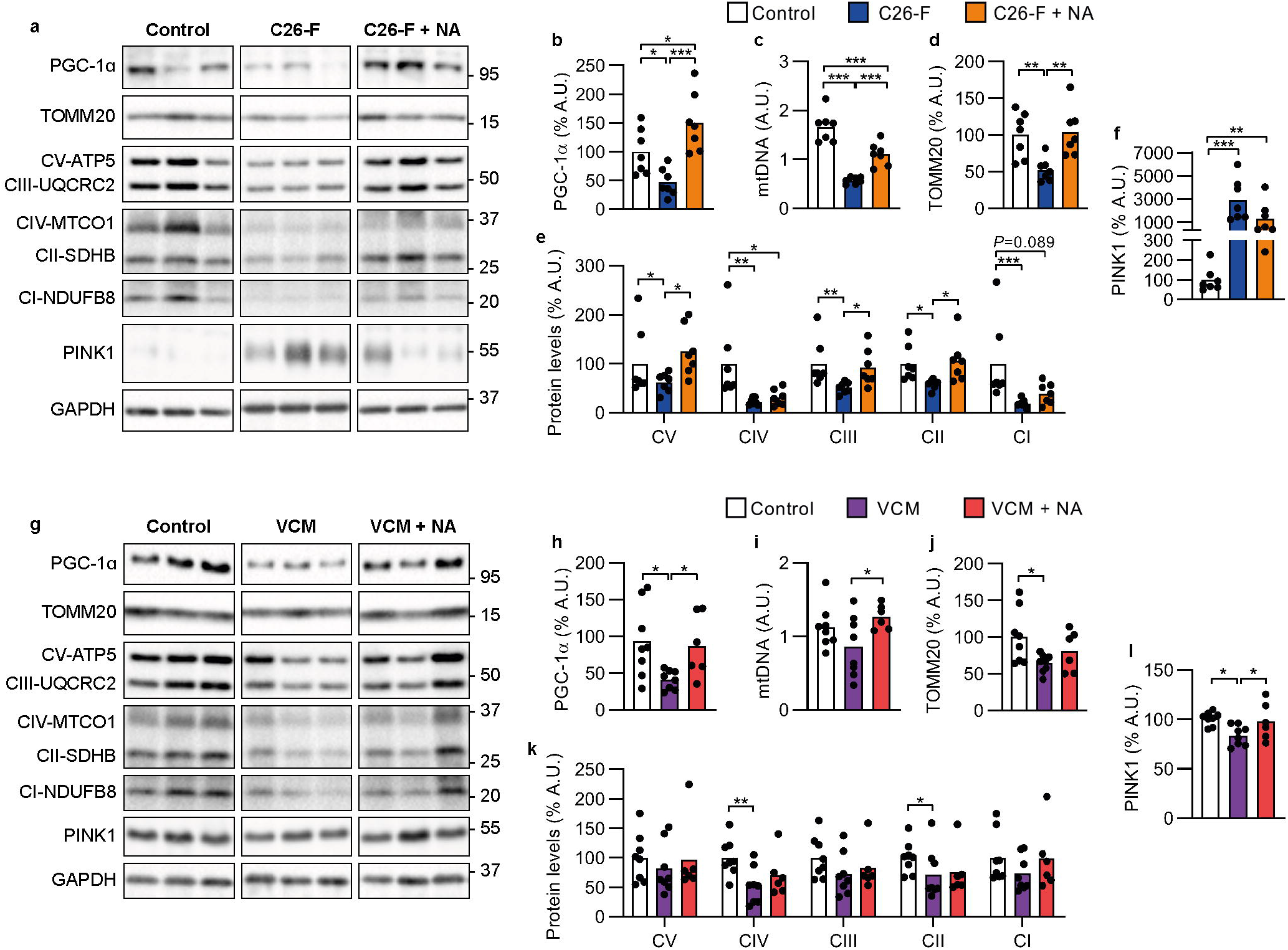
Impact of NA on skeletal muscle mitochondrial respiratory capacity and markers of mitochondrial biogenesis in CC mouse models. **a-f** Assays run on *gastrocnemius* (GSN) muscle from control (*n*=6-7), C26-F (*n*=7) and C26-F + NA (*n*=7) groups: **a** representative western blotting bands for PGC-1α, TOMM20, OXPHOS complex subunits (ATP5, UQCRC2, MTCO1, SDHB and NDUF88), PINK1 and GAPDH; **b** expression of PGC-1α protein as assessed by densitometry analysis of western blotting bands. GAPDH protein expression was used as loading control; **c** mtDNA amount presented as a ratio of mtDNA genome to nuclear DNA genome; **d-f** Expression of TOMM20 protein (**d**), OXPHOS proteins (**e**) and PINK1 (**f**) as assessed by densitometry analysis of western blotting bands (**a**). GAPDH protein expression was used as loading control. **g-l** Assays run on GSN muscle from control (*n*=8), VCM (*n*=8) and VCM + NA (*n*=5-6) groups: **g** representative western blotting bands for PGC-1α, TOMM20, OXPHOS complex subunits, PINK1 and GAPDH; **h** expression of PGC-1α protein as assessed by densitometry analysis of western blotting bands (**g**). GAPDH protein expression was used as loading control; **i** mtDNA amount presented as a ratio of mtDNA genome to nuclear DNA genome; **j-l** Expression of TOMM20 protein (**j**), OXPHOS proteins (**k**) and PINK1 (**l**) as assessed by densitometry analysis of western blotting bands. GAPDH protein expression was used as loading control. Data are shown as means with individual values. Statistical analysis was performed with ANOVA + Fisher’s LSD for normally distributed data and with Kruskal-Wallis + Uncorrected Dunn’s test for non-normal data. *P<0.05, **P<0.01 and ***P<0.001. NA; niacin, A.U.; arbitrary units.

Similarly to the C26-F model, the skeletal muscle of VCM mice showed a robust decrease in PGC-1α protein levels (Fig. 4g,h) and significant reductions in TOMM20 and the OXPHOS complex subunits MTCO1 (CIV) and SDHB (CII) (Fig. 4g,j-k), although no significant changes in mtDNA abundance were observed when compared to controls (Fig. 4i). Yet no accumulation of PINK1 was observed in the muscle of VCM mice (Fig. 4g,l). NA rescued PGC-1α levels (Fig. 4g,h), increased mtDNA content (Fig. 4i), and partially improved TOMM20 and OXPHOS complex subunit MTCO1 (CIV) and NDUF88 (CI) (Fig. 4g,j-k). As in C26-F mice, VCM mice showed downregulation of genes promoting mitochondrial biogenesis, although in this case, NA partially corrected the expression of *Erra* (Supplementary Fig. 2p). Overall, NA treatment ameliorates skeletal muscle mitochondrial status in two distinct models of experimental CC.

### Niacin corrects liver NAD^+^ deficiency and partially improves hepatic mitochondrial alterations

CC is a complex metabolic disease where the liver has a crucial role in the control of systemic energy and glucose metabolism^2^. As the liver also contributes to the systemic regulation of NAD^+^ synthesis and recycling^21^, we examined how CC and NA treatment influence liver condition and NAD^+^ metabolism in C26F and VCM mice.

C26F mice showed liver hypertrophy with depleted hepatic glycogen and total glutathione levels (Supplementary Fig. 3a-d), together with a severe decline in blood glucose levels (Supplementary Fig. 3e). NA supplementation slightly improved total glutathione content and glycemia whereas NA had no significant effects on liver size or hepatic glycogen levels (Supplementary Fig. 3a-e). All hepatic NAD metabolites (NAD^+^, NADH, NADP^+^ and NADPH) were dramatically reduced in C26-F mice as compared to controls (Fig. 5a). NAD^+^ depletion likely originated from a strong downregulation of NAD^+^ biosynthetic enzymes of salvage and Preiss-Handler pathways including the liver isoform of nicotinamide riboside kinase, *Nrk1* (Supplementary Fig. 3f), not from the enhanced NAD^+^ consumption via PARPs (Supplementary Fig. 3g). NA restored all hepatic NAD metabolite concentrations in C26-F mice (Fig. 5a). The livers of C26-F mice showed increased protein synthesis as opposed to skeletal muscle, even with the accumulation of autophagosomes (LC3B-II). Neither protein synthesis nor LC3B levels were influenced by NA (Supplementary Fig. 3h-j). Although no changes in liver mtDNA amount were detected (Fig. 5b), a decline in the protein expression of TOMM20 and OXPHOS complex subunits was observed in VCM compared to controls (Fig. 5c,d). Similarly to skeletal muscle, a reduction of transcripts was observed for the activators of mitochondrial biogenesis *Tfam* and *Erra* in C26-F mice (Supplementary Fig. 3k). Interestingly, NA-treatment increased mtDNA amount above control levels and partially rescued the expression of TOMM20 and the OXPHOS II, III and IV complex subunits (Fig. 5b-d). No transcriptional induction of mitochondrial biogenesis markers occurred in the liver after NA supplementation (Supplementary Fig. 3k).

**Figure 5.**
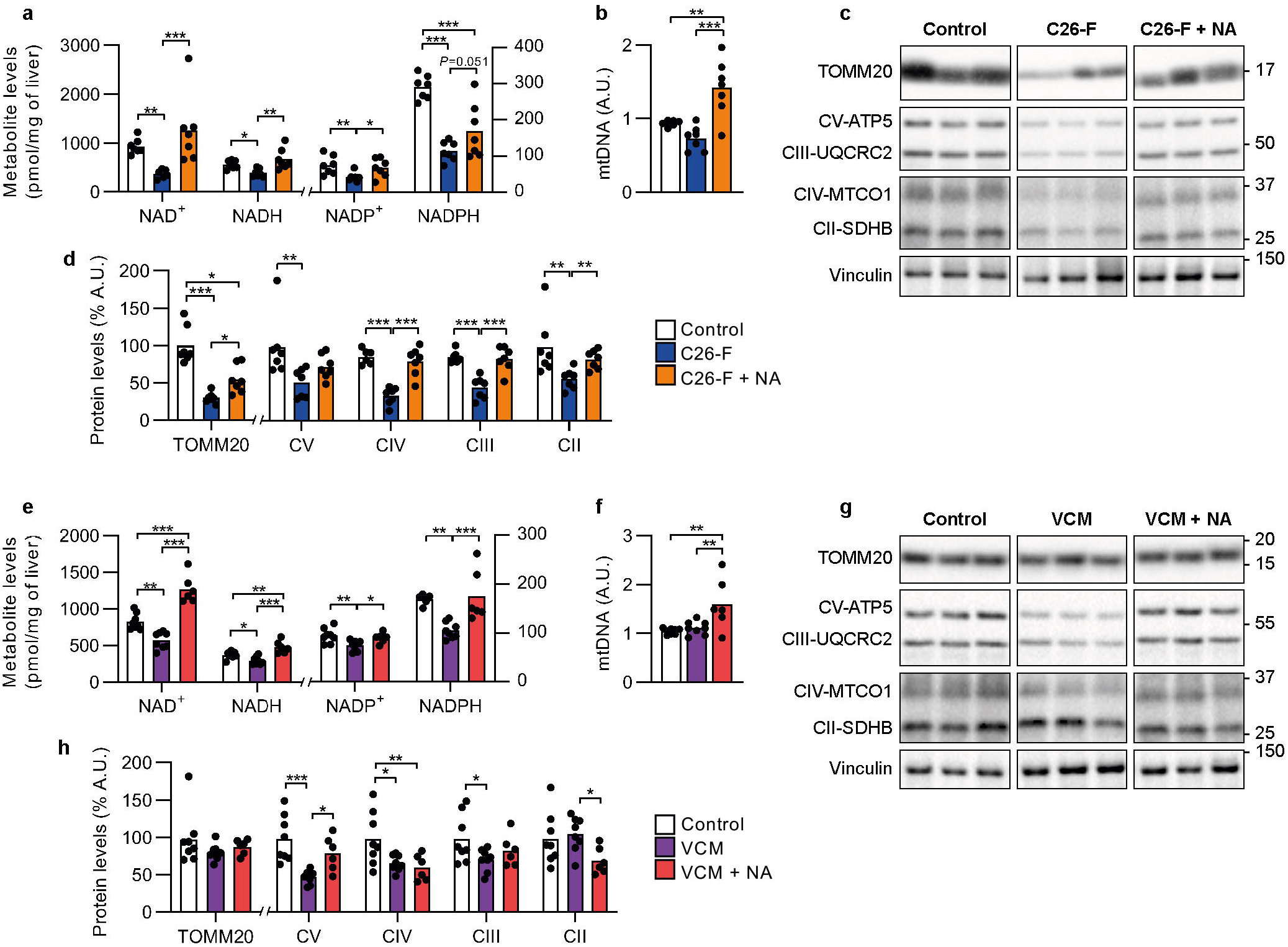
Impact of NA on hepatic NAD^+^ and mitochondrial metabolism in CC mouse models. **a-d** Assays run on frozen liver from from control (*n*=6-7), C26-F (*n*=7) and C26-F + NA (*n*=7) groups: **a** hepatic NAD^+^ levels as pmol normalized to liver weight; **b** mtDNA amount presented as a ratio of mtDNA genome to nuclear DNA genome; **c** Representative western blotting bands of TOMM20, OXPHOS complex subunits (ATP5, UQCRC2, MTCO1, SDHB and NDUF88) and Vinculin; **d** Expression of TOMM20 and OXPHOS proteins as assessed by densitometry analysis of western blotting bands (**c**). Vinculin protein expression was used as loading control. **e-h** Assays run on frozen liver from control (*n*=6-8), VCM (*n*=8) and VCM + NA (*n*=6) groups: **e** hepatic NAD^+^ levels as pmol normalized to liver weight; **f** mtDNA amount presented as a ratio of mtDNA genome to nuclear DNA genome; **g** Representative western blotting bands of TOMM20, OXPHOS complex subunits (ATP5, UQCRC2, MTCO1, SDHB and NDUF88) and Vinculin; **h** Expression of TOMM20 and OXPHOS proteins as assessed by densitometry analysis of western blotting bands (**g**). Vinculin protein expression was used as loading control. Data are shown as means with individual values. Statistical analysis was performed with ANOVA + Fisher’s LSD for normally distributed data and with Kruskal-Wallis + Uncorrected Dunn’s test for non-normal data. \**P*<0.05, \*\**P*<0.01 and \*\*\**P*<0.001. NA; niacin, A.U.; arbitrary units.

As in C26-F mice, VCM mice showed hepatomegaly (Supplementary Fig. 3l) and a dramatic reduction in hepatic NAD^+^, NADH, NADP^+^ and NADPH content as compared to controls (Fig. 5e). The expression of NAD^+^ biosynthetic genes and PARP activity remained fairly stable (Supplementary Fig. 3m,n) in the liver. NA supplementation, not affecting liver size, restored hepatic NAD metabolite concentrations in VCM mice (Fig. 5e, Supplementary Fig. 3l). Hepatic mtDNA amount and protein expression of TOMM20 did not differ between VCM and control mice (Fig. 5f-h). Yet protein expression of OXPHOS complex III, IV and V subunits were significantly decreased in tumor-bearing animals (Fig. 5g,h). NA supplementation did not influence TOMM20 expression but it increased mtDNA amount and ATP5 (CV subunit) expression as compared to non-treated VCM mice (Fig. 5f-h). The transcription of mitochondrial biogenesis markers partially decreased in VCM mice while NA only improved the expression of *Tfam* (Supplementary Fig. 3o). In conclusion, these findings reveal that CC is characterized by the deficiency of hepatic NAD metabolites and mitochondrial abnormalities that are partially restored boosting NAD^+^ metabolism with NA.

## DISCUSSION

Disturbed skeletal muscle NAD^+^ metabolism has recently emerged as a molecular determinant of murine CC^4^. Our study reveals that the downregulation of muscle NAD^+^ biosynthetic enzyme *NRK2* is a common feature of murine and human CC, allowing to identify patients with metabolic disturbances. Importantly, rescuing NAD^+^ levels protects from cancer- and chemotherapy-induced muscle wasting in mice.

NAD^+^ depletion and perturbed NAD^+^ biosynthesis are well established pathophysiological factors of diseases characterized by muscle mitochondrial dysfunction and disturbed energy metabolism, such as mitochondrial myopathies and sarcopenia^8,13^. As summarized in figure 6, muscle NAD^+^ depletion occurs mainly in severe CC mouse models. In contrast, the downregulation of *Nrk2* was detected in the skeletal muscle in all mouse models including the milder and chronic VCM model of CC, not presenting with NAD^+^ depletion. This finding suggests that *Nrk2* loss may precede the development of NAD^+^ metabolism disturbances and muscle loss. Consistently, we show for the first time that cancer patients exhibit muscle *NRK2* repression (Fig. 6), already in pre-cachectic state and exacerbated in overt cachexia. As muscle *NRK2* loss occurred independently from CC status, *i*.*e*. body weight loss and/or sarcopenia, this emphasizes that NAD^+^ metabolism could be a viable target for early interventions to improve cancer patient health before overt or refractory CC ensue. In addition, our results highlight the inability of the present CC assessment procedures to detect muscle or systemic metabolic abnormalities that strongly impair cancer patient outcome and quality of life.

**Figure 6.**
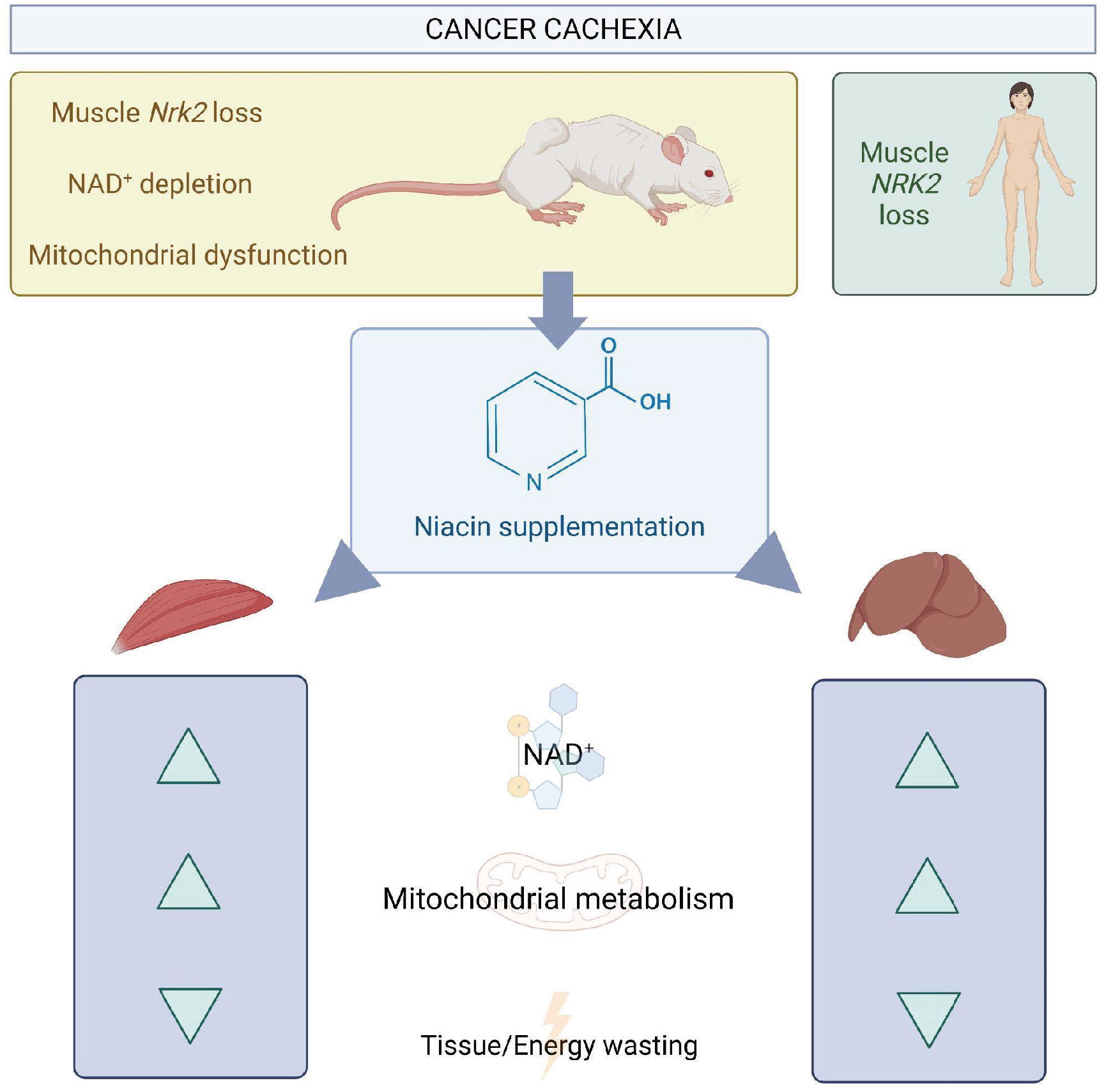
Graphical representation of the main finding of this study. Observations in cancer patients confirm the data obtained in three distinct murine models of experimental cancer cachexia and are supported by positive results in the intervention study involving severe (C26-F) and chronic (VCM) cancer cachexia. The beneficial effects produced by niacin on NAD content, mitochondrial homeostasis and energy metabolism are generalized despite model specific alterations are present.

Previous mouse studies have demonstrated that *Nrk2* plays a redundant role in basal muscle NAD^+^ biosynthesis^22,23^. In contrast, *Nrk2* is typically upregulated during metabolic energy stress and NAD^+^ deficiency to support NAD^+^ production^23–25^. This is contrary to our findings of consistent *Nrk2* downregulation in CC, suggesting that either the skeletal muscle has impaired adaptation to NAD^+^ deficiency or that *Nrk2* loss plays a primary role in determining the altered NAD^+^ and energy metabolism in CC. In line with the latter notion, low muscle *NRK2* expression was associated with metabolite alterations in both skeletal muscle and serum of cancer patients, highlighting the future possibility to set up simple and fast venous blood sampling to diagnose energy metabolism disturbances in CC patients. Overall, our results indicate that muscle NRK2 loss is a common feature of murine and human CC and that NRK2 might have a disease-specific role in the regulation of energy homeostasis.

Beneficial effects of increasing intracellular NAD^+^ levels have been demonstrated in various muscle and metabolic diseases^13,26–29^. In agreement with these previous studies, NA partially rescued the depleted NAD^+^ metabolites in the skeletal muscle, counteracted muscle wasting and improved muscle function and protein synthesis in C26-F mice. Although the VCM mice showed no depletion of muscle NAD metabolites at baseline, NA restored muscle mass and normalized autophagic markers. These improvements possibly originate, especially in C26-F mice, from a better maintenance of mitochondrial energy metabolism rather than NA having a direct effect on protein metabolism. Consequently, the energized muscles have a lesser need for muscle degradation to fulfill systemic energy demands. In both CC models, NA likely increased mitochondrial biogenesis via a PGC-1α-mediated mechanism. A similar positive effect on muscle mitochondrial metabolism has been observed upon NAD^+^ boosting therapies in several rodent studies^7^. In addition, in a recent human study from our group, NA improved muscle mitochondrial biogenesis and muscle strength in healthy individuals and patients with mitochondrial myopathy^13^. Collectively, our murine study indicates that NA has a therapeutic potency on CC regardless of the muscle NAD^+^ content, that may vary according to the severity of CC and/or the exposure to chemotherapy.

Considering that the liver performs a wide range of energetically demanding processes, it has been suggested that hepatic metabolism requires proper NAD^+^ homeostasis^30–32^. In line, dysregulated mitochondrial, lipid and glucose metabolism are associated with hepatic NAD^+^ deficiency^33,34^. Here we provide the first evidence that both severe and mild CC mouse models exhibit pronounced hepatic NAD metabolite depletion. The underlying cause for hepatic NAD^+^ deficiency may be related to perturbed NAD^+^ biosynthesis, at least in C26-F mice. These findings revealed that NAD^+^ metabolism aberrations are rather of systemic nature in CC than specifically distinctive for skeletal muscle. Despite NA effectively rescued hepatic NAD metabolite levels and muscle wasting in both models, with only a partial restoration effect on liver mitochondrial metabolism, a causal relationship cannot be established between liver dysfunction and muscle wasting in the currently adopted murine CC models, suggesting that the beneficial impact of NA on skeletal muscle is not likely secondary to the rescue of liver metabolism.

In conclusion, our findings encourage investigating NRK2-targeted therapeutic options to improve disturbed energy metabolism in CC. In addition, the results demonstrate that NA has a therapeutic effect on both cancer- and chemotherapy-induced cachexia in mice. The effectiveness of NA in variable experimental conditions that reflect the broad human spectrum of CC increases the translational value of our findings. Although deeper insight on NAD^+^ metabolism impairments in different conditions of human CC are still required, our study highlights the necessity of NAD^+^ to support energy metabolism in CC and paves the way for the development of novel vitamin B3-based therapies to effectively target the multifaceted aspects of CC.

## MATERIALS AND METHODS

All the reagents used in this work were obtained from Merck-MilliporeSigma (St. Louis, MO, USA) unless differently indicated.

### Animals and experimental design

Experimental animals were cared for in compliance with the Italian Ministry of Health Guidelines and the Policy on Humane Care and Use of Laboratory Animals (NRC, 2011). The experimental protocols were approved by the Bioethical Committee of the University of Torino (Torino, Italy) and the Italian Ministry of Health (Aut. Nr. 579/2018-PR). The animals were maintained on a regular dark-light cycle of 12:12 hours with controlled temperature (18-23°C) and free access to food and water during the whole experimental period. B6.Cg-Tg(Vil1-cre)997Gum/J (Villin-Cre) and B6.Cg-Msh2^tm2.1Rak^/J (Msh2^loxP^), mice were purchased from The Jackson Laboratory (Bar Harbor, CA, USA) and were crossed to obtain the Villin-Cre/Msh2^loxP/loxP^(VCM) offspring, leading to the conditional knock-out of the Msh2 gene in the enterocytes of the intestinal mucosa, accelerating the formation of intestinal adenomas/adenocarcinomas^15^. The presence of each transgenic construct was assessed through Melt Curve Analysis (RT-qPCR) using the following primers: Villin-Cre (forward: 5’-TTCTCCTCTAGGCTCGTCCA-3’ and reverse: 5’-CATGTCCATCAGGTTCTTGC-3’) and Msh2^loxP^ (wild-type: 5’-GATGATGTGTGAAGCCTGCAT-3’, mutant: 5’-CCTCTTGAGGGGAATTGAAGT-3’ and common: 5’-AGGTTAAAAACCAGAGCCTCAACT-3’).

Two distinct experiments were performed in this study:

Animal experiment 1: 21 6-month-old wild-type BALB/c mice weighing approximately 20g (Charles River, Wilmington, MA) were divided into 3 groups (*n*=7): healthy controls, C26-F and C26-F+NA. Females were used to avoid the fighting characteristic among male cagemates subjected to severe cachexia protocols. C26-F mice were subcutaneously inoculated with 5 × 10^5^ Colon26 (C26) carcinoma cells on the back and treated with Folfox chemotherapy (6 mg/Kg oxaliplatin, 25 mg/Kg 5-fluorouracil, 90 mg/Kg leucovorin) at days 7, 14 and 21 after tumor inoculation. Mice of the C26-F + NA group were administered a daily dose of NA (150 mg/kg dissolved in tap water) by gavage. Healthy controls received an equal saline injection excluding the cell inoculum and were daily treated with tap water. Grasping strength was assessed on day 0 and the day after every Folfox administration (days 8, 15 and 22). Oral treatments with NA started 4 days after tumor injection. At day 28 post-C26 implantation, mice were injected with an intraperitoneal dose of 40 μmol/Kg puromycin 30 min prior to euthanasia in order to assess the relative rate of protein synthesis (SUnSET methodology)^35^. The amount of puromycin incorporated into nascent peptides was detected by western blotting, using a specific anti-puromycin antibody.

Animal experiment 2: one-year-old male VCM mice (*n*=6) were treated with a daily dose of NA (150 mg/kg, up to 4.5 mg/day) by gavage for 28 days. Age and gender-matched VCM mice (*n*=8) and *Msh2*^loxP/loxP^ mice (*n*=8) were used as non-treated tumor-bearers and healthy controls, respectively.

Muscle samples from KPC mice derivate from a previous animal experiment^14^. Briefly, 8-week-old wild-type C57BL/6 male mice were subcutaneously inoculated with 0.7 × 10^6^ cells KPC cells (*n*=6) and animals were terminated 5 weeks after tumor implantation. Healthy age-matched C57BL/6 mice were used as controls (*n*=6).

In all experimental protocols, body weight and food intake were monitored every other day, and the animals were daily examined for signs of distress. At the endpoint, the mice were anesthetized with 2% isoflurane in O2, blood was collected by cardiac puncture and euthanasia was performed by means of cervical dislocation. Several tissues were excised, weighed, frozen in liquid nitrogen and stored at -80°C for further analyses.

### Collection of human skeletal muscle and serum samples

The human samples originate, with some additions, from a previous study^16^. From 2015 to 2020 we enrolled consecutive patients with colorectal or pancreatic cancer and control patients undergoing surgery for benign diseases at the 3rd Surgical Clinic of the University Hospital of Padova. The research project was approved by the Ethical Committee for Clinical Experimentation of Padova (protocol number 3674/AO/15). All patients joined the protocol according to the guidelines of the Declaration of Helsinki and the written informed consent was obtained from participants. The muscle biopsy was performed at the time of the planned surgery by a cold section of a *rectus abdominal* fragment (1×0.5 cm) immediately frozen and conserved in liquid nitrogen for gene expression analysis and metabolome profiling. Serum samples were obtained from blood samples retrieved prior to any surgical manipulation. Demographics and clinical data, including medications and comorbidities noted as having potential confounding effects on skeletal muscle homeostasis^16^ were collected from all patients (Supplementary Table 1). Cancer patients were classified as cachectic in cases of >5% weight loss in the 6 months preceding surgery, >2% weight loss with either body mass index (BMI) <20 or low muscle mass defined by the skeletal muscle index (SMI) cut-offs described by Martin *et al*^36^. SMI values were quantified using the preoperative CT scans as previously described^16^. Based on gene expression analysis, we selected 10 cancer patients with the highest (comparable to healthy controls) and 10 with the lowest (almost 10-fold decrease) *NRK2* levels (Supplementary Table 2, 3). We performed metabolome profiling of muscle and serum samples in these two subgroups.

### Metabolome analysis

About 10 mg of skeletal muscle or 50 μl of serum were used for metabolite extraction and analysis. The samples were flash frozen upon collection and sent for further processing to the Metabolomics Expertise Center, VIB Center for Cancer Biology, KULeuven Department of Oncology, Leuven, Belgium. The extraction was performed adding 99 or 19 volumes (for muscles or sera, respectively) of 80% methanol, containing 2 uM d27 myristic acid as internal standard. The mixture was centrifuged at 20.000 x g for 15 min at 4°C to precipitate proteins and insoluble material, the supernatant transferred to a fresh new tube. 10 μl of each sample was loaded into a Dionex UltiMate 3000 LC System (Thermo Scientific Bremen, Germany) equipped with a C-18 column (Acquity UPLC -HSS T3 1. 8 μm; 2.1 × 150 mm, Waters) coupled to a Q Exactive Orbitrap mass spectrometer (Thermo Scientific) operating in negative ion mode. A step gradient was carried out using solvent A (10 mM TBA and 15 mM acetic acid) and solvent B (100% methanol). The gradient started with 5% of solvent B and 95% solvent A and remained at 5% B until 2 min post injection. A linear gradient to 37% B was carried out until 7 min and increased to 41% until 14 min. Between 14 and 26 minutes the gradient increased to 95% of B and remained at 95% B for 4 minutes. At 30 min the gradient returned to 5% B. The chromatography was stopped at 40 min. The flow was kept constant at 0.25 mL/min at the column was placed at 40°C throughout the analysis. The MS operated in full scan mode (m/z range: [70.0000-1050.0000]) using a spray voltage of 4.80 kV,capillary temperature of 300 °C, sheath gas at 40.0, auxiliary gas at 10.0. The AGC target was set at 3.0E+006 using a resolution of 140000, with a maximum IT fill time of 512 ms. Data collection was performed using the Xcalibur software (Thermo Scientific). The data were obtained by integrating the peak areas (El-Maven – Polly - Elucidata). Data analysis was performed using the free online resource https://www.metaboanalyst.ca/ version 5.0.

### Liver glycogen and glutathione content

Liver glycogen concentration was assessed using a commercially available system (MAK016 Glycogen assay Kit). Briefly, liver fragments of about 50 mg were cold homogenized in water (10% w/vol) with a bead homogenizer (Bullet Blender, New Advance, Troy, NY, USA), boiled for 5’ and centrifuged for 5’ at 13,000 x *g*. The supernatant was collected and diluted 100-fold before adding 10 μl to a 96 well plate. The assay was performed following manufacturer’s instructions and using a glycogen titration curve in order to extrapolate quantitative data.

Glutathione was determined as previously described^37^, with slight modifications^38^. Briefly, liver fragments of about 50 mg were cold homogenized in water (10% w/vol) with a bead homogenizer, deproteinized on ice using 5% metaphosphoric acid and centrifuged at 15,000 x g for 2 min. The supernatants were treated with 4 M triethanolamine to reach pH 7.4. GSH concentration was determined after 2 min incubation with 5,50-dithiobis-2-nitrobenzoic acid (DTNB) by measuring the production of 50-thio-2-nitrobenzoic acid (TNB) at 412 nm on a 96-well microplate reader. Suitable volumes of diluted glutathione reductase (6 U/mL) and of NADPH (4 mg/mL) were then added to evaluate total glutathione level (GSH + GSSG). GSSG content was calculated by subtracting GSH content from total glutathione levels.

### Assessment of NAD metabolite levels

NAD^+^, NADH and the phosphorylated metabolites NADP^+^ and NADPH were measured from pulverized *gastrocnemius* (GSN) muscle and liver with a slightly modified conventional colorimetric method^4^ (for further information, see https://www.nadmed.fi/). Metabolite levels were normalized to tissue mass used for analysis or the total protein content of the sample.

### PARP activity

PARP activity was analyzed from pulverized liver and *gastrocnemius* (GSN) muscle utilizing HT Colorimetric PARP/Apoptosis Assay Kit (R&D Systems, Minneapolis, MN, USA) according to manufacturer’s instructions (n=6-8 per group). Data were normalized with the protein amount of the samples.

### Mitochondrial DNA amount quantification

Total DNA, including mitochondrial DNA, was extracted from approximately 10 mg of pulverized skeletal muscle and approximately 3 mg of pulverized liver from C26-F and VCM mice with the standard phenol-chloroform method followed by ethanol precipitation. The amount of mtDNA was determined as the ratio of mitochondrial rRNAs, 16s and cytochrome c oxidase subunit II (Cox2) genomic regions, to the geometric mean of nuclear uncoupling protein 2 (Ucp2) and hexokinase-2 (Hk2) genomic regions using RT-qPCR. Primer sequences are listed in Supplementary Table X. RT-qPCR was carried out in triplicates with 2 ng of template DNA per well using Maxima SYBR Green qPCR Master Mix (Thermofisher Scientific) and the CFX Connect Real-Time PCR Detection System (Bio-Rad). Data analysis was conducted with standard curve method with qBASE+ software (Biogazelle).

### RNA isolation and RT-qPCR analysis

Approximately 30 mg of GSN muscle and liver were lysed and processed to isolate high-quality RNA using the standard phenol-chloroform method. RNA concentration was quantified by means of spectrophotometry. Total RNA was retro-transcribed using a cDNA synthesis kit (Bio-Rad, Hercules, CA, USA or Qiagen, Hilden, Germany) and transcript levels were determined by RT-qPCR using the SsoAdvanced SYBR Green Supermix (Bio-Rad) or Maxima SYBR Green qPCR Master Mix (Thermofisher Scientific) and the CFX Connect Real-Time PCR Detection System (Bio-Rad) with 10 ng of cDNA per well. Every RT-qPCR was validated by analyzing the respective melting curve and run in parallel to no reverse transcriptase control (NRT) to exclude potential artifacts from genomic DNA contamination. Gene expression was normalized to the geometric mean of housekeeping gene expression and represented as relative expression according to primer efficiency assessed using serial dilutions of pooled samples (standard curve method). Data analysis was conducted in Microsoft Excel and qBASE+ software (Biogazelle). As for human muscle biopsies, total RNA was extracted from approximately 20 mg of rectus abdominal muscle using TRIzol (Thermo Fisher Scientific). 1 ug of RNA was reverse transcribed using the SuperScript IV Reverse Transcriptase (Thermo Fisher Scientific). Gene expression was analyzed by qRT-PCR using the PowerUp SYBR Green Master Mix (Applied Biosystems). Data were normalized to *Actb* gene expression. Primer sequences are listed in Supplementary Table 4.

### Western blotting

Approximately 50 mg of GSN muscle and liver were mechanically homogenized using bead homogenizer in RIPA buffer (50 mM Tris-HCl pH 8.0, 5 mM EDTA pH 8.0, 1% Igepal CA-630, 0.5% sodium deoxycholate, 0.1% SDS) containing protease inhibitors (0.5 mM PMSF, 0.5 mM DTT, 2 μg/ml leupeptin, 2 μg/ml aprotinin) and phosphatase inhibitors (P0044). Next, homogenates were sonicated for 10 s at low intensity, centrifuged at 15000 *g* for 5 min at 4°C and the supernatant was collected. Total protein concentration was quantified with Bradford reagent (Bio-Rad) using BSA as protein concentration standard. Equal amounts of protein (10-30 μg) were heat-denatured (except when assessing OXPHOS expression) in sample-loading buffer (50 mM Tris-HCl, pH 6.8, 100 mM DTT, 2% SDS, 0.1% bromophenol blue, 10% glycerol), resolved by SDS-PAGE electrophoresis (4561086, Bio-Rad) and transferred to nitrocellulose membranes (1704159, Bio-Rad). The filters were blocked with 5% nonfat dry milk in Tris-buffered saline containing 0.05% Tween (TBS-Tween) and then incubated overnight with antibodies directed against specific proteins: AMPK (07-181, Millipore), p-AMPK (#2535, Cell Signaling), GAPDH (G8795), LC3B (L7543), NRK2 (produced in Dr. Gareth G Lavery’s lab), OXPHOS Antibody Cocktail (ab110413, Abcam), PGC-1α (ab3242, Merck Millipore), PINK1 (SAB2500794), Puromycin (EQ0001, Kerafast), TOMM20 (ab186735, Abcam) and Vinculin (sc73614, Santa Cruz Biotechnology). Peroxidase conjugated IgGs (Bio-Rad) were used as secondary antibodies. Three 5 min washes with TBS-Tween were performed after each antibody incubation. After incubation with Clarity Western ECL substrate (170-5061, Bio-Rad), bands were developed using the ChemiDoc XRS+ imaging system (Bio-Rad). Densitometric analysis on the obtained images was performed using the Image Lab software (Bio-Rad).

### Data representation and statistics

Data are presented using bar (mean) and dot plots (individual values) unless differently stated. Data representation and statistical tests were performed with Prism (version 9, GraphPad) software. Outliers were identified using ROUT (Q=1%) and excluded from the analysis. The normality of distributions was evaluated by the Shapiro-Wilk test. Unless differently stated in the figure legend, the significance of the differences was evaluated by appropriate two-sided statistical tests: Student’s “t”-test or analysis of variance (ANOVA) for normal distribution and Mann–Whitney test or Kruskal–Wallis test for non-normal distribution. ANOVA was followed by the Fisher’s Least Significant Difference (LSD) test, whereas the Kruskal–Wallis test was followed by the Uncorrected Dunn’s test to assess differences of planned comparisons among groups.

## AUTHOR CONTRIBUTION

Conceptualization, M.B., N.P., J.J.H., E.P. and F.P.; Formal Analysis, M.B. and N.P.; Investigation, M.B., N.P., C.F., K.T., M.Y.H. and R.S.; Resources, S.Z., L.M., P.E.P., R.K., M.S. and R.S.; Writing – Original Draft, M.B., N.P., E.P. and F.P.; Writing – Review & Editing, S.Z., P.E.P., M.S., J.J.H., and R.S.; Supervision, E.P. and F.P.; Funding Acquisition, E.P. and F.P.

## ACKNOWLEDGMENTS

We are grateful to Dr. Gareth G Lavery (Department of Biosciences, Nottingham Trent University) who kindly donated the antibody against the NRK2 protein. Also, we thank Valentina Audrito (Department of Medical Sciences, University of Turin) for reviewing and advising on the first draft of the manuscript. This work was supported by Fondazione AIRC (IG 2018—ID. 21963 project, PI: F.P.), the Finnish Cancer Foundation and Finnish Cancer Center FICAN South (PIs: E.P. and Dr Tommi Järvinen, respectively), and by two post-doctoral Fellowships from Fondazione Umberto Veronesi (ID2496 and ID3519 to R.S).

