## Supplementary Figure and Tables for "NAD^+^ repletion with niacin counteracts cancer cachexia"

### Supplementary Fig. 1

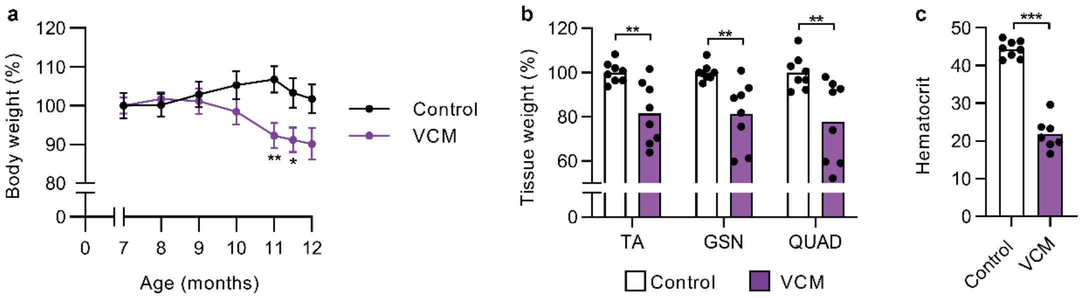


**Supplementary Fig. 1. Body weight loss, muscle mass depletion and reduced hematocrit of 12-month-old tumor-bearing VCM mice.**

**a** Body weight change of control and tumor-bearing VCM mice (months 7-11 n=7-8) between 7 and 12 months of age (months 7-11: control and VCM *n*=8; months 11.5-12: control *n*=7, VCM *n*=5). **b** Wet weight of *tibialis anterior* (TA), *gastrocnemius* (GSN) and *quadriceps femoris* (QUAD) muscles represented as a percentage of the mean of the control group. **c** Hematocrit of 12-month-old control and VCM mice. Data display: **a** are means ± SEM; **b-c** are means with individual values. Statistical analysis was performed using Student's t-test, *$P$<0.05, **$P$<0.01 and ***$P$<0.001 (control *n*=8; VCM mice *n*=7-8).

#### Supplementary Fig. 2

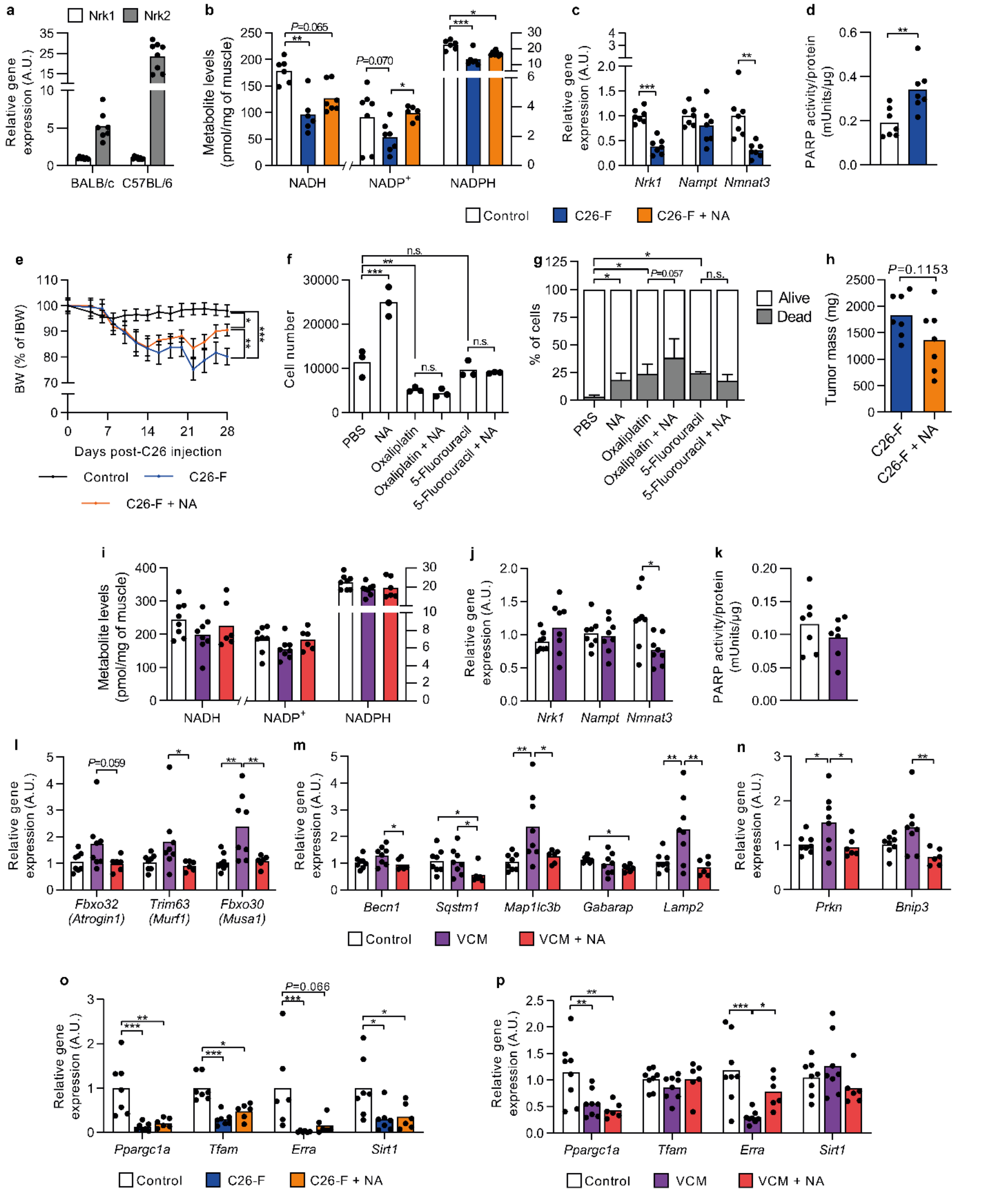


**Supplementary Fig. 2. In the skeletal muscle, niacin replenishes NAD metabolites in C26-F mice and reduces the expression of atrogenes in VCM mice.**

**a** Relative expression of *Nrk1* and *Nrk2* genes in the skeletal muscle of BALB/c (*n*=7) and C57BL/6 (*n*=8) mouse strands. **b** Levels of NADH, NADP+ and NADPH as pmol normalized to muscle mass (*n*=6-7). **c** Relative expression of genes involved in NAD+ biosynthesis in the muscle of Control and C26-F groups (*n*=6-7). **d** Total PARP activity as mUnits normalized to total protein content. **e** Body weight change of control, C26-F and C26-F + NA groups upon C26 tumor implantation (*n*=7). Data are represented as percentage of mouse initial body weight (IBW). **f-g** Assessment of cell number (**f**) and death ratio of C26 cells (**g**) after 24h of exposure to 1mM NA, 10 μM oxaliplatin and 1 μM 5-fluorouracil alone, and both chemotherapeutics combined individually with NA (*n*=3 independent wells). **h** Tumor mass in C26-F and C26F + NA groups after animal euthanasia (*n*=7). **i** Levels of NADH, $NADP^+$ and NADPH as pmol normalized to muscle mass (*n*=6-8). **j** Relative expression of genes involved in $NAD^+$ biosynthesis in the skeletal muscle (*n*=8). **k** Total PARP activity as mUnits normalized to total protein content (*n*=7). **l-n** Relative expression of E3 ubiquitin ligases (**l**), autophagy (**m**) and mitophagy (**n**) genes in the skeletetal muscle (*n*=5-8). **o, p** Relative expression of genes involved in mitochondrial biogenesis (**o**) C26-F (*n*=6-7) and (**q**) VCM (*n*=6-8) experimental models, respectively. Data display: **a-d, f, h-p** are means with individual values; **e, g** are means ± SEM. Statistical analysis was performed either with Student's t-test or ANOVA + Fisher's LSD for normally distributed data and with Kruskal-Wallis + Uncorrected Dunn's test for non-normal data. *$P<0.05$, **$P<0.01$ and ***$P<0.001$. NA; niacin, A.U.; arbitrary units.

#### Supplementary Fig. 3

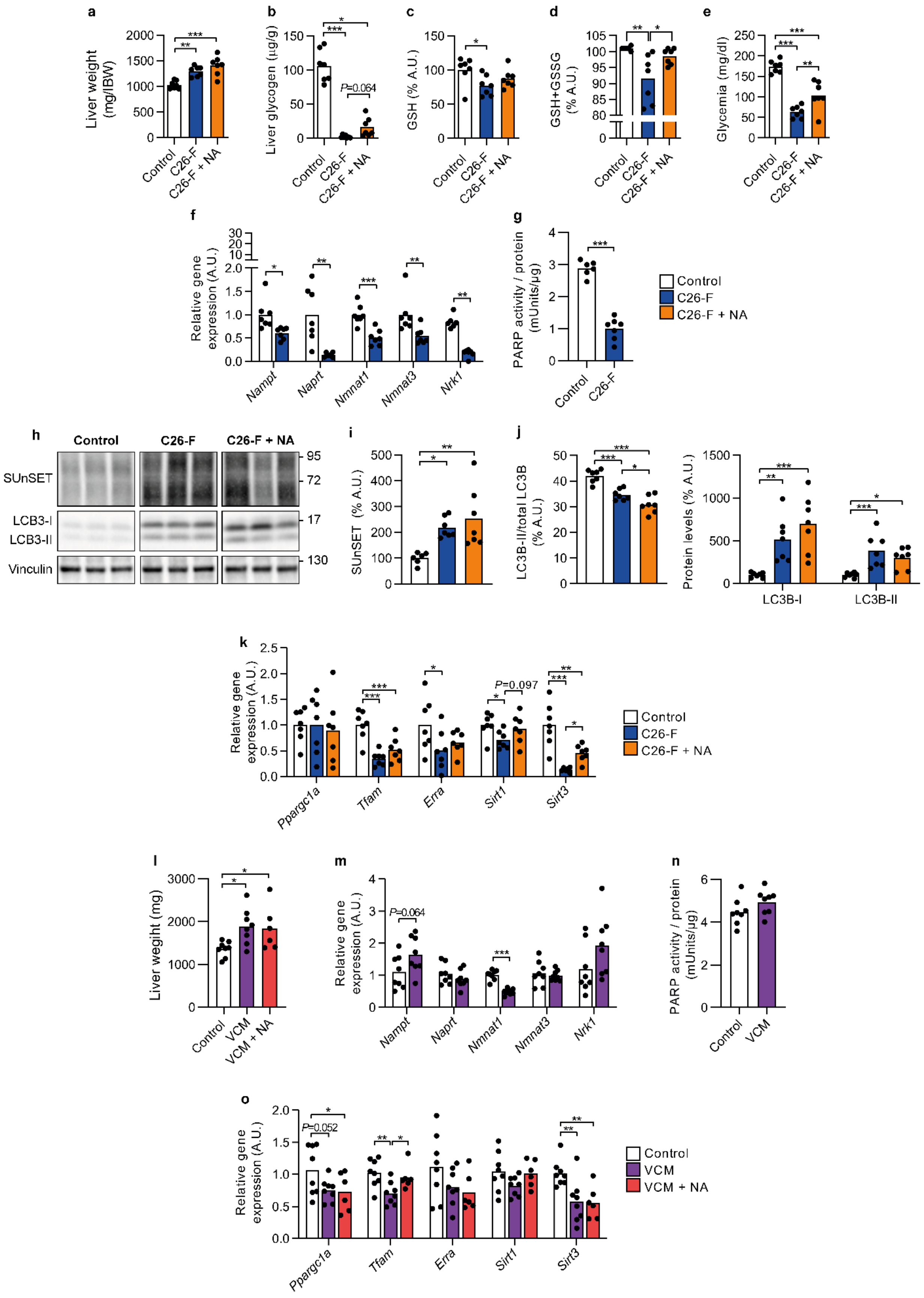

**Supplementary Fig. 3. Alterations in NAD$^+$ biosynthesis, protein metabolism and mitochondrial homeostasis in the liver of C26-F and VCM mice are only mildly influenced by niacin treatment.**

**a** Liver wet weight normalized by initial body weight (IBW) from control, C26-F and C26-F + NA groups (*n*=7). **b** Hepatic glycogen content (*n*=7). **c, d** Content of reduced (**c**) and total (**d**) hepatic glutathione (*n*=6-7). **e** Glycemia calculated as mg of glucose per dL of blood (*n*=7). **f** Relative expression of genes involved in NAD$^+$ biosynthesis in the liver (*n*=6-7). **g** Total PARP activity as mUnits normalized to total protein content (*n*=6-7). **h-j** Representative western blotting bands (**h**) and densitometry analysis of puromycin incorporation (SUnSET analysis) (**i**) and LC3B (**j**) proteins (*n*=6-7). Vinculin protein expression was used as loading control. **k** Relative expression of genes involved in mitochondrial biogenesis (*n*=6-7). **l** Liver wet weight from control, VCM and VCM + NA groups (*n*=6-8). **m** Relative expression of genes involved in NAD biosynthesis in the liver (*n*=8). **n** Total PARP activity as mUnits normalized to total protein content (*n*=8). **o** Relative expression of genes involved in mitochondrial biogenesis (*n*=6-8). All bar plots display means with individual values. Statistical analysis was performed either with Student’s t test or ANOVA + Fisher’s LSD for normally distributed data and with Mann-Whitney or Kruskal-Wallis + Uncorrected Dunn’s test for non-normal data. *P<0.05, **P<0.01 and ***P<0.001. NA; niacin, A.U.; arbitrary units.

**Supplementary Table 1. Patient population's characteristics**

| | Control (n=28) | PC (n=49) | C (n=53) | Total (n=130) | P value |
|---|---|---|---|---|---|
| F % (n) | 53.6 (15) | 44.9 (22) | 50.9 (27) | 49.2 (64) | ns |
| M % (n) | 46.4 (13) | 55.1 (27) | 49.1 (26) | 50.8 (66) | |
| Age, mean years ± SD (Min-Max) | 62.7 ± 12.9 (40-86) | 68.0 ± 13.3 (40-88) | 68.0 ± 8.9 (49-88) | | 0.04 Cntr vs PC |
| BMI (kg/m$^2$), mean ± SD | 25.1 ± 3.3[a] | 25.4 ± 4.1 | 23.1 ± 3.6 | | 0.01 (Cntr vs C); 0.002 (PC vs C) |
| BMI category (kg/m$^2$) | | | | | 0.02 |
| < 20, % (n) | 0 (0) | 6.1 (3) | 22.6 (12) | | |
| 20,0 to 24,9, % (n) | 59.3 (16) | 46.9 (23) | 52.8 (28) | | |
| 25,0 to 29,9, % (n) | 33.3 (9) | 32.7 (16) | 20.8 (11) | | |
| ≥ 30 | 7.4 (2) | 14.3 (7) | 3.8 (2) | | |
| Weight loss (%), mean ± SD (Min/Max) | -0.7 ± 2.7 (0/-13)[a] | -0.4 ± 1.1 (0/-4.7) | -11.3 ± 5.5 (-3.1/-25) | | <0,0001 (Cntr vs C ; PC vs C) |
| Drugs, % (n) | 50 (11)[b] | 55.1 (27) | 56.6 (30) | | ns |
| Co-morbidities, % (n) | 13.6 (3)[b] | 22.4 (11) | 28.3 (15) | | ns |

All demographic and clinical data were collected at the time of surgery. Differences between groups were analyzed by one-way ANOVA (continuous variable that passed the normality test) or Kruskal-Wallis test (continuous variable that didn't pass the normality test) with Benjamini, Krieger and Yekutieli adjustment and $\chi^2$ test (categorical variables).

Abbreviations: F, female; M, male; BMI, body mass index calculated as patient weight (kg)/height (m)$^2$. Weight loss in 6 months before biopsies were calculated with the following formula: [(current weight [kg] - weight 6 months ago [kg])/weight 6 months ago (kg)] X 100; negative values indicate weight loss.

[a] Control patients with BMI and weight loss information: n=27

[b] Control patients with information on current medication and co-morbidities: n=22

**Supplementary Table 2. Metabolomic patient subset's characteristics**

| | Healthy (n=10) | High NRK2 (n=10) | Low NRK2 (n=10) | Total (n=30) | P value |
|---|---|---|---|---|---|
| F % (n) | 50 (5) | 50 (5) | 50 (5) | 50 (15) | ns |
| M % (n) | 50 (5) | 50 (5) | 50 (5) | 50 (15) | |
| Age, mean years ± SD (Min-Max) | 64.8 ± 15.2 (40-81) | 71.5 ± 8.8 (61-86) | 64.1 ± 9.3 (49-76) | | ns |
| BMI (kg/m$^2$), mean ± SD | 25.2 ± 2,5 | 22.3 ± 2.6 | 22.8 ± 3.1 | | 0.04 (Healthy vs High); 0.08 (Healthy vs Low) |
| BMI category (kg/m$^2$) | | | | | ns |
| < 20, % (n) | 0 (0) | 20 (2) | 30 (3) | | |
| 20,0 to 24,9, % (n) | 60 (6) | 70 (7) | 40 (4) | | |
| 25,0 to 29,9, % (n) | 40 (4) | 10 (1) | 30 (3) | | |
| ≥ 30 | 0 (0) | 0 (0) | 0 (0) | | |
| Weight loss (%), mean ± SD (Min/Max) | 0 ± 0 (0/0) | -7.6 ± 10.1 (0/-25) | -10.1 ± 5.9 (0/-17.5) | | 0.04 (Healthy vs High); 0.0008 (Healthy vs Low) |
| Drugs, % (n) | 70 (7) | 40 (4) | 10 (1) | | 0.02 |
| Co-morbidities, % (n) | 10 (1) | 10 (1) | 0 (0) | | ns |

All demographic and clinical data were collected at the time of surgery. Differences between groups were analyzed by one-way ANOVA (continuous variable that passed the normality test) or Kruskal-Wallis test (continuous variable that didn't pass the normality test) with Benjamini, Krieger and Yekutieli adjustment and $\chi^2$ test (categorical variables).

Abbreviations: F, female; M, male; BMI, body mass index calculated as patient weight (kg)/height (m)$^2$. Weight loss in 6 months before biopsies were calculated with the following formula: [(current weight [kg] - weight 6 months ago [kg])/weight 6 months ago (kg)] X 100; negative values indicate weight loss.

**Supplementary Table 3. Metabolomic oncological patient subset's characteristics**

| | High NRK2 (n=10) | Low NRK2 (n=10) | P value |
|---|---|---|---|
| Cachectic, % (n) | 40 (4) | 80 (8) | ns |
| Low muscle mass, % (n) | 70 (7) | 50 (5) | ns |
| Myosteatosis, % (n) | 50 (5) | 30 (3) | ns |
| Tumor type, % (n) | | | |
| CRC | 20 (2) | 0 (0) | 0.04 |
| Pancreas | 80 (8) | 60 (6) | |
| Other gastrointestinal | 0 (0) | 40 (4) | |
| Tumor stage, % (n) | | | ns |
| 0 | 10 (1) | 10 (1) | |
| I | 0 (0) | 10 (1) | |
| II | 30 (3) | 20 (2) | |
| III | 30 (3) | 40 (4) | |
| IV | 30 (3) | 20 (2) | |
| CT < 4 weeks, % (n) | 10 (1) | 0(0) | ns |

All clinical data were collected at the time of biopsy.

CT scans for body composition analysis were collected within 60 days before surgery.

Cancer patients are classified having low muscle mass in case of SMI<41 $cm^2/m^2$ in females; SMI<43 $cm^2/m^2$ if BMI <25, and SMI<53 $cm^2/m^2$ if BMI ≥25 in males and having myosteatosis in case of HU <33 if BMI ≥25, and <41 if BMI <25, regardless the sex (Martin L. *et al.*). Skeletal muscle index was calculated as lumbar total muscle cross-sectional area ($cm^2$)/height $(m)^2$. Differences between groups were analyzed by $\chi^2$ test of Fisher's exact test (categorical variables).

Abbreviations: CRC, colorectal cancer. CT, chemotherapy exposure within 4 weeks prior to muscle biopsy. Other gastrointestinal: duodenum, ampulla of Vater, distal bile duct

**Supplementary Table 4. Primers used for qPCR analyses**

| Species | Gene | Forward | Reverse |
|---|---|---|---|
| mouse | *36b4* | GGCCCTGCACTCTCGCTTTC | TGCCAGGACGCGCTTGT |
| mouse | *Becn1* | TGAAATCAATGCTGCCTGGG | CCAGAACAGTATAACGGCAACTCC |
| mouse | *Beta-actin* | CTGGCTCCTAGCACCATGAAGAT | GGTGGACAGTGAGGCCAGGAT |
| mouse | *Bnip3* | GTCGCCTGGCCTCAGAAC | CCCATTGCCATTGCTGAAGT |
| mouse | *Erra* | ACTGCCACTGCAGGATGAG | CACAGCCTCAGCATCTTCAA |
| mouse | *Fbxo30/Musa1* | TCGTGGAATGGTAATCTTGC | CCTCCCGTTTCTCTATCACG |
| mouse | *Fbxo32/Atrogin1* | GCAAACACTGCCACATTCTCTC | CTTGAGGGGAAAGTGAGACG |
| mouse | *Gabarap* | TCCGTGCTGAAGATGCCTTG | TCTTCTTCATGGTGTTCCTGGTA |
| mouse | *H2bc4* | TACAACAAGCGCTCGACCA | TCTGCTCCTCTTGGCAGG |
| mouse | *Hprt* | GAGGAGTCCTGTTGATGTTGCCAG | GGCTGGCCTATAGGCTCATAGTGC |
| mouse | *Lamp2* | GCTGAACAACAGCCAAATTA | CTGAGCCATTAGCCAAATACAT |
| mouse | *Map1lc3b* | CACTGCTCTGTCTTGTGTA | TCGTTGTGCCTTTATTAGTG |
| mouse | *Nampt* | GAACAGATACTGTGGCGGGA | CCAAGCCGTTATGGTACTGTG |
| mouse | *Naprt* | GCAGGACTGTATGCGCTTTCT | GAAGCGGCACACCAGGGA |
| mouse | *Nmnat1* | TGTGCCCAAGGTGAAATTGCT | CCACGATTTGCGTGATGTCC |
| mouse | *Nmnat3* | CACGAATATGCACCTGCGCT | CATTGACGGGTGAGATGATGC |
| mouse | *Nrk1* | CCCAACTGCAGCGTCATATC | CCTTGAGCACTTTCCAAGGC |
| mouse | *Nrk2* | CACCTCAGGACCAGTCACCT | CTGTTGGTCAGGGTGGTCTT |
| mouse | *Ppargc1a (1)* | AAGTGTGGAACTCTCTGGAACTG | GGGTTATCTTGGTTGGCTTTATG |
| mouse | *Ppargc1a (2)* | GCAACATGCTCAAGCCAAAC | TGCAGTTCCAGAGAGTTCCA |
| mouse | *Prkn* | CGTGTGATTTTTGCCGGGAAG | GGTCCACTCGTGTCAAGCTC |
| mouse | *Sirt1* | GTCTCCTGTGGGATTCCTGA | ACACAGAGACGGCTGGAACT |
| mouse | *Sirt3* | CTGAAACCGGATGGCGTTTG | ACCATGACCACCACCCTACT |
| mouse | *Sqstm1* | GGCCACCTCTCTGATAGCTTCT | GACATTGGGATCTTCTGGTGGA |
| mouse | *Tfam* | AAGTGTTTTTCCAGCATGGG | GGCTGCAATTTTCCTAACCA |
| mouse | *Trim63/Murf1* | GGGCCATTGACTTTGGGACA | TCTCCTTCTTCATTGGTGTTCTTCT |
| human | *ACTB* | GGGAAATCGTGCGTGACA | GGACTCCATGCCCAGGA |
| human | *NRK2* | AGGATGACTTCTTCAAGCCCC | GGGCACGGGCAAACTTCT |
